# Augmenting Transcriptome Annotations through the Lens of Splicing Evolution

**DOI:** 10.1101/2024.11.04.621892

**Authors:** Xiaofei Carl Zang, Ke Chen, Irtesam Mahmud Khan, Mingfu Shao

## Abstract

Alternative splicing (AS) is a ubiquitous mechanism in eukaryotes. It is estimated that 90% of human genes are alternatively spliced. Despite enormous efforts, transcriptome annotations remain, nevertheless, incomplete. Conventional means of annotation were largely driven by experimental data such as RNA-seq and protein sequences, while little insight was shed on understanding transcriptomes and alternative splicings from the perspective of evolution. This study addresses this critical gap by presenting TENNIS (Transcript EvolutioN for New Isoform Splicing), an evolution-based model to predict unannotated isoforms and refine existing annotations without requiring additional data. The model of TENNIS is based on two minimal premises–AS isoforms evolve sequentially from existing isoforms, and each evolutionary step involves a single AS event. We formulate the identification of missing transcripts as an optimization problem and parsimoniously find the minimal number of novel transcripts. Our analysis showed approximately 80% of multi-transcript groups from six transcriptome annotations satisfy our evolutionary model. At a high confidence level, 40% of isoforms predicted by TENNIS were validated by deep long-read RNA-seq. In a simulated incomplete annotation scenario, TENNIS dramatically outperforms two randomized baseline approaches by a 2.25-3 fold-change in precision or a 3.5-3.9 fold-change in recall, after controlling the same level of recall or precision of the baseline methods. These results demonstrate that TENNIS effectively identifies missing transcripts by complying with minimal propositions, offering a powerful approach for transcriptome augmentations through the lens of alternative splicing evolutions. TENNIS is freely available at https://github.com/Shao-Group/tennis.

## 1 Introduction

Alternative splicing (AS) is a ubiquitous and prevalent mechanism in eukaryotes. It alternatively splice-in or splice-out some exons from the same pre-mRNA [30]. AS increases the diversity of transcript isoforms [5,35]. AS happens more frequently and more independently than previously estimated [38]. It is estimated that over 90% of human genes are alternatively spliced [20,33]. There are four basic types of AS events [30]: (1) cassette exon (CE), also known as exon skipping/inclusion, (2) alternative 3’ splicing site (A3), (3) alternative 5’ splice sites (A5), and (4) intron retention (IR) (Fig. 1A). Some complex AS events, such as multiple exon skipping or exclusive exons, can be considered as the synergy of two basic AS events.

**Fig. 1:**
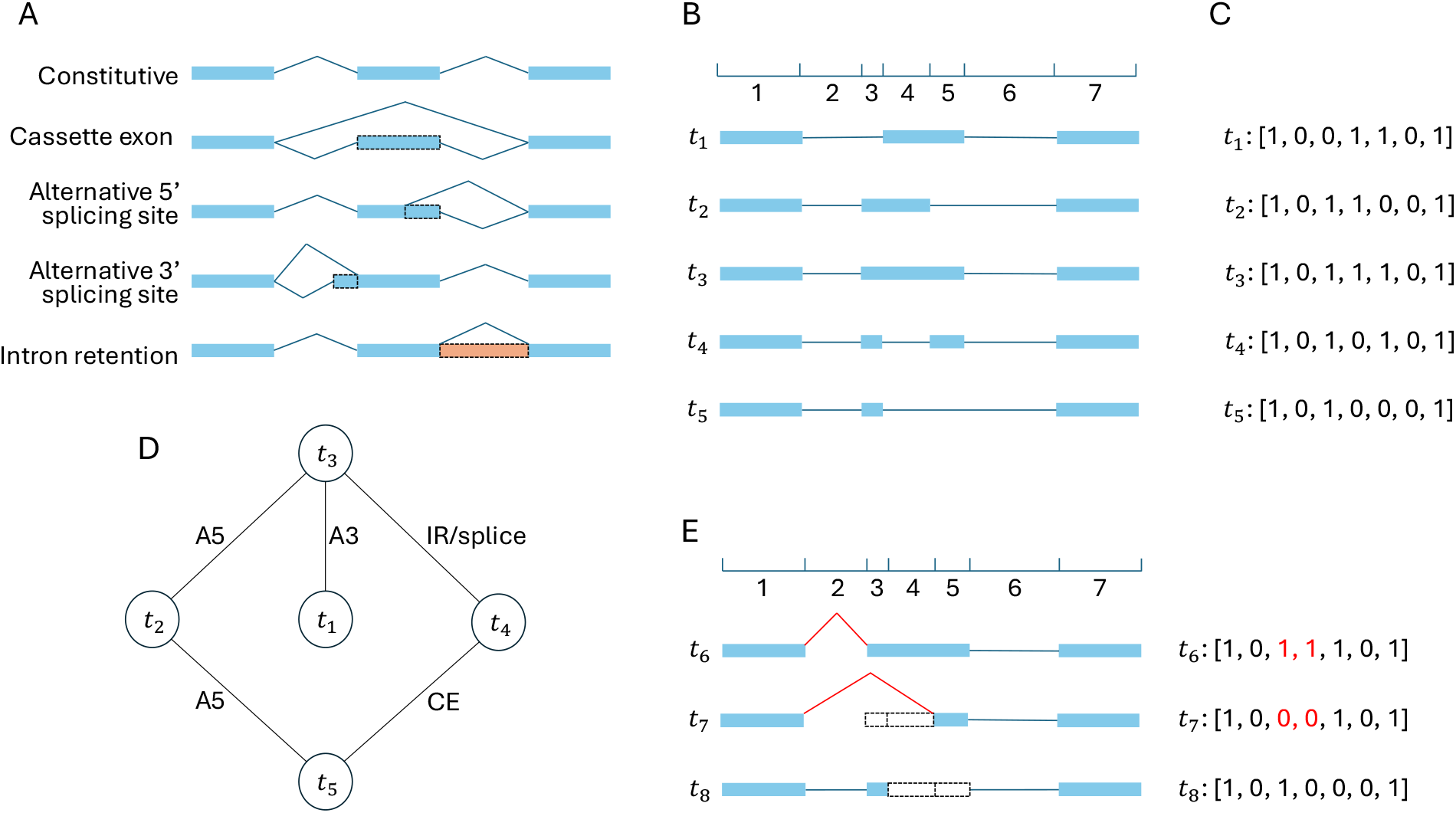
(A) Four basic alternative splicing types. Blue rectangle: exon; Peach rectangle: retained intron; Dashed rectangle: alternative (partial) exon/intron; Blue polyline: splice junction. (B) Splice sites of all transcripts divide the genome into several sub-regions. This example shows a group of 5 transcripts (*t*_1_, *t*_2_, *t*_3_, *t*_4_, *t*_5_) divides the genome into 7 regions. (C) Each transcript is encoded as a binary vector indicating which regions are spliced in (1) or spliced out (0). This example encodes panel B transcripts as vectors of length 7. (D) Potential AS events between panel B transcripts. Only exactly one AS event is permitted in between. (E) Skipping multiple consecutive partial exons is one AS event. The red splice junctions and binary bits illustrate that converting *t*_6_ to *t*_7_ is one A3 event, but two consecutive partial exons are skipped. Similarly, converting *t*_6_ to *t*_8_ is one A5 event (junctions not shown). CE: cassette exon; A5: alternative 5’ splice sites; A3: alternative 3’ splicing site; IR: intron retention.

The study of AS is extensive, ranging from the mechanisms of splicing regulation [34,6] to the functions of splicing isoforms and their associations with diseases [31,7,27]. One important angle of studying AS is through evolution. It is known that AS is under rapid evolution and is elastically shaped by environments [39]. Elucidating the evolutionary relationship across splicing isoforms originated from the same pre-mRNA is crucial, as it is closely related to functional diversification of genes and offers a powerful tool to study splicing regulation [10,28]. For example, AS might have originated through DNA mutations in the splicing sites, control sequences, and the evolution of splicing regulators [9,3]. It was also reported that multi-intron genes may precede the emergence of AS, and in primate species, AS events combine independently with each other so that novel AS isoforms emerge [3,38]. Despite these biological advances, there remains a significant shortage of mathematical models that quantitatively characterize splicing evolution.

The catalog of all splicing isoforms of all genes, i.e. transcripts, for a species is called the transcriptome. These transcripts not only transcribe genetic information to encode proteins but also play important regulatory and functional roles [29,16]. Various biological and biomedical studies are heavily dependent on fine-grained transcriptome annotations, including the quantification of transcripts, the curation of a single-cell expression atlas, the identification of aberrant splicing in disease-related samples, and comparative transcriptomics.

Over the past decades, tremendous effort has been put into constructing and improving the annotations of transcriptomes, especially the model organisms. For illustration, the major consortia for the annotation of the human species include RefSeq [14], Ensembl [1], CHESS [22], and MANE [17]. These annotations were primarily conducted in a data-driven manner, where one of the classic ways is to perform assembly from RNA-seq data [25,23]. The assemblies are often additionally augmented or validated by experimental data. For example, NCBI annotations, including RefSeq, also consider transcript sequences, reads in the SRA database, CAGE-Seq, amino acid sequences, and curated data from other sources [14,18]. The Ensembl annotation consolidates information from cDNAs, protein sequences, RNA-seq, and manual curations [1]. CHESS is based on a large-scale RNA-seq of nearly ten thousand samples [22]. The MANE annotation constitutes a consensus between RefSeq and ENSEMBL with manual curations [17]. Despite the significant amount of computational tools, pipelines, and manual curations, the transcriptome annotations are not complete even for model organisms [25,37,36]. Humans, the undoubtedly most-studied species, had a continually increasing number of recorded genes and transcripts from GRCh37 to GRCh38 [26], to T2T-CHM13 [19]. Annotation for other model organisms, mouse or Drosophila, are also incomplete, as novel transcripts were found with higher sequencing depth and more comprehensive sequencing experiments [12,32,2].

In this work, we propose a mathematical model for splicing evolution. Based on this model, we develop a tool called TENNIS (Transcript EvolutioN for New Isoform Splicing) that is able to predict missing isoforms in an annotation (without using any external sequencing data). Our model characterizes the AS evolution trajectory based on two simple premises. First, evolution does not create new splicing isoforms out of thin air, rather, it modifies and adapts existing ones; and second, evolution takes baby steps, namely, each isoform is derived from its predecessor through a single AS event. Under this model, AS isoforms in each group (precise definition in Section 2.1) should form a connected graph where vertices are the isoforms, and edges represent a single basic splicing event (CE, A3, A5, or IR), see Fig. 1D for an example. We validate this model with the transcriptome annotations of several model organisms, including human, mouse, Drosophila, zebrafish, maize, and Arabidopsis. We found that this model can explain approximately 80%-90% of the transcript groups for six transcriptome annotations that we investigated.

If the AS isoforms in a group cannot be represented as an evolutionary graph defined above, i.e., it does not satisfy our model, then this is evidence that some transcripts might be missed in the annotation. We develop a computational approach to determine potentially missing isoforms for such cases. We formulate it as an optimization problem following the parsimony principle: to seek the minimum number of missing transcripts whose inclusion connects all the AS isoforms. We develop an exact algorithm to solve this problem. Specifically, we devise a new satisfiability (SAT) formulation to determine if adding *k* missing transcripts suffices, for each *k* = 1, 2, 3, …, and hence the smallest such *k* gives the optimal solution. The resulting SAT instances can be solved with existing solvers such as Glucose [4]. We apply TENNIS to the annotations of model species. The majority of the transcript groups that do not satisfy our model miss only one isoform. Across different filtering strategies, 30%-40% of the predicted novel transcripts were supported by deep long-read RNA-seq studies, substantially higher than a random baseline prediction. To further validate our model and TENNIS, we simulated an incomplete annotation scenario by removing some isoforms and found TENNIS can retrieve 25.4%-41.3% missing isoforms with a 28%-40.6% precision. TENNIS has nearly 2.25-3 times of the precision or 3.5-3.9 times of the recall of two randomized baseline approaches after controlling the same level of recall or precision.

## 2 Methods

### 2.1 An evolution-inspired model

TENNIS models the evolution of alternative splicing (AS) within a transcript group (defined below) based on two premises: (1) AS isoforms evolve sequentially, with each isoform being derived from a predecessor; and (2) each isoform must originate from its parent through a single AS event (CE, A3, A5, or IR) per evolutionary step. The rationale behind the second premise is that AS events arise independently through mutations in splicing sites or regulatory elements and it is less likely to have two mutations occur simultaneously [3,38]. Consequently, all isoforms of a transcript group should be connected via single AS events. If not, then it indicates that isoform(s) are missing from the annotation or lost function and therefore are not present in the current annotation.

The framework of TENNIS is as follows. It takes a transcriptome or an assembly, i.e., a set of annotated or assembled transcripts in gtf format, as input. It first partitions all transcripts into transcript groups (defined below). Within each group, it constructs the evolutionary relationship using a graph, determines evidence of missing isoforms, and if evidence presents, identifies the missing isoforms.

We focused on analyzing AS isoforms originating from the same pre-mRNA. That is, TENNIS groups transcripts that share the same alternative transcription start site (TSS) and alternative transcription termination site (TTS) together, referred to as a “transcript group”. We denote by *𝒯*_*S*_ the set of transcript groups with just a single transcript, and by *𝒯*_*M*_ the set of transcript groups with two ore more transcripts. Fig. 1B shows an example of a transcript group with 5 transcripts. Although TSS and TTS are two events that also produce diverse transcripts, the pre-mRNAs are already different for transcripts with such events [15]. Hence, the AS processes are more different between transcripts with alternative TSS or TTS [2,24].

Next, TENNIS builds a graph for each transcript group. In the graph, the collection of nodes represents all isoforms and the collection of edges represents that two isoforms are convertible via a single AS event. For example, Fig. 1D illustrates the graph for transcripts in Fig. 1B. Details of the construction of graphs are described in Section 2.2. We say that a transcript group does not present evidence of missing isoform(s) if the graph is connected (i.e., the group consists of a single connected component). Otherwise, TENNIS recruits a minimal number of additional nodes to make all components connected. These reconstructed nodes/transcripts are regarded as missing isoforms. This step is modeled as an optimization problem and solved by transforming it into a satisfiability (SAT) formulation, detailed in Section 2.3.

### 2.2 Constructing evolution trajectory and identifying missing isoforms

Let *T* be a transcript group. TENNIS encodes each isoform in *T* as a binary vector, depicting exonic or intronic regions. First, genomic coordinates of all splicing sites of all isoforms in *T* are collected and then the genome is split into smaller regions according to those coordinates (Fig. 1B). Let *n* be the number of resulting genomic regions. Clearly, each exon or intron spans either one region or several consecutive regions. Hence, every isoform in *T* can be described using indices of spliced-in regions (i.e. exonic regions) and indices of spliced-out regions (i.e. intronic regions). Therefore, by encoding the exonic region as 1 and intronic regions as 0, an isoform is encoded as a length-*n* binary vector. For example, a 1 at position *i* of the binary vector means the *i*-th genomic region is covered by an exon in this isoform and vice versa (see Fig. 1C). Assume *T* contains *m* isoforms. Then *T* can be represented as an *m × n* binary matrix, denoted as *M*.

The benefits of binary encoding of isoforms are that, besides clarity and conciseness, all simple AS events can be represented as the flip of a bit or several consecutive bits. While an exon is split into multiple smaller regions (in this case, called partial-exons) due to A3/A5 events in another isoform, this exon is accordingly coded as multiple 1’s. Partial-introns are defined likewise. Hence, the A3/A5 event can be represented as a flip of the bits of those corresponding partial-exons to partial-introns. CE and IR are also represented as 1-to-0 or 0-to-1 flips. In another way, all simple AS events can be considered as the inclusion or exclusion of one or multiple consecutive regions.

Given an *m×n* binary matrix *M* representing a transcript group, a graph will be constructed. Each annotated isoform is denoted as a vertex. An edge may be added between two vertices if their isoforms can convert to each other by one AS event, *i*.*e*. flip of consecutive 0’s to 1’s or consecutive 0’s to 1’s. Edges are undirected, since 0-to-1 and 1-to-0 flips are symmetric. This also reflects the invertible property of the basic events.

Providing no missing transcript, all vertices should be in one connected component. In this case, we say that transcript group *T* satisfies our evolutionary model, and call *T* a transcript group in 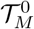. Otherwise, one or more isoforms are said to be missed in *T*. It is important to note that, due to the minimality of our model, neither direction of the reasoning is decisive, that is, it is possible that *T* misses some isoforms but the resulting graph remains connected, and it is also possible that, the graph is not connected but *T* does not miss any unannotated isoform.

In the case that the graph contains more than one connected component, TENNIS will reconstruct missing isoforms. We formulate this task as an optimization problem, that is, to find a minimum number of isoforms such that adding them to *T* will result in a graph with just one connected component. We design an algorithm, termed TENNIS-SAT (*M, k*), described in Section 2.3, that takes matrix *M* and an integer *k ≥* 1 as input, and answers if adding *k* isoforms suffices to make the resulting graph connected, and if yes, also returns the binary representation of the *k* additional isoforms. Using TENNIS-SAT as subroutine, starting with *k* = 1, TENNIS employs an iterative approach, that calls TENNIS-SAT(*M, k*) in each iteration and increases *k*, until either the subroutine returns yes (and the *k* isoforms) or a maximum iteration number is reached. As a compromise of computational time and accuracy, the default maximum iterations, which is also the maximum number of missing isoforms that TENNIS attempts to reconstruct, is 4. According to our experiments, with this threshold, the model can explain more than 97% of the investigated transcript groups (Table 1). Transcript group *T* will be assigned to a category 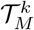, if TENNIS determines that a minimum of *k* transcripts are missed in *T, k* = 1, 2, 3, 4. *T* will be assigned to category 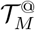 if the maximum iteration reached, which means *T* misses more than 4 transcripts, or TENNIS fails to finish in 15 minutes.

**Table 1:** Summary statistics of the number of transcript groups in each category.

| Species | Annotation | $\mathcal{T}_M^0$ | $\mathcal{T}_M^1$ | $\mathcal{T}_M^2$ | $\mathcal{T}_M^3$ | $\mathcal{T}_M^4$ | $\mathcal{T}_M^\oplus$ | $\mathcal{T}_M$ | $\mathcal{T}_S$ |
| --- | --- | --- | --- | --- | --- | --- | --- | --- | --- |
| Human/hg38 | GENCODE | 11960(62%) | 4277(22%) | 1654(9%) | 676(3%) | 322(2%) | 536(3%) | 19425 | 125777(87%) |
| Human/hg38 | RefSeq | 20852(78%) | 3951(15%) | 1169(4%) | 435(2%) | 199(1%) | 278(1%) | 26884 | 63541(70%) |
| Mouse | GRCm39 | 17178(84%) | 2321(11%) | 599(3%) | 190(1%) | 92(0%) | 102(0%) | 20482 | 49524(71%) |
| Drosophila | dm6 | 2433(78%) | 451(14%) | 116(4%) | 46(1%) | 28(1%) | 42(1%) | 3116 | 17938(85%) |
| Zebrafish | GRCz11 | 7455(88%) | 673(8%) | 155(2%) | 62(1%) | 21(0%) | 61(1%) | 8427 | 41836(83%) |
| Maize | NAM-5.0 | 16799(88%) | 1612(8%) | 332(2%) | 142(1%) | 53(0%) | 46(0%) | 18984 | 83398(81%) |
| Arabidopsis | TAIR10 | 1481(88%) | 147(9%) | 33(2%) | 10(1%) | 2(0%) | 2(0%) | 1675 | 9105(84%) |

It is common that multiple optimal solutions exist. This means, for a transcript group *T* in 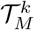, different sets of *k* isoforms may make the resulting graph connected. For example, in Fig. 1D, if both *t*_2_ and *t*_4_ were missing, then either of them would be an optimal solution of size 1. TENNIS is able to return all optimal solutions. This offers an additional critical signal to decide if a constructed missing isoform is correct or not. The intuition is, that if there are multiple optimal solutions, and an isoform appears in all of them, then it is more likely to be truly missed than these just appear in one solution. We, therefore, for each reconstructed isoform in the union of all optimal solutions, introduce a measure “Percentage In (PctIn)”, defined as the number of solutions containing this isoform divided by the total number of solutions. In the above example, both *t*_2_ and *t*_4_ will be in the output with a PctIn value of 0.5.

### 2.3 TENNIS-SAT

Given an *m × n* binary matrix *M* representing all isoforms in a transcript group *T*, and an integer *k* representing the maximum number of missing isoforms allowed to be added, we use a SAT formulation to decide if adding *k* isoforms is sufficient to connect the graph. Similar to existing isoforms, the unknown novel isoforms are represented as a vector of binary variables. For simplicity, they are appended to *M*, and all isoforms are represented by rows in *M*. That means, *M*_*i*_ is a length-*n* vector of known binary values for *i* = 1, *· · ·, m*, while *M*_*i*_ is a length-*n* vector of unkown binary variables for *i* = *m* + 1, *· · ·, m* + *k*.

Since the aim is to construct a connected graph, the presence of a spanning tree in the graph is necessary and sufficient. The spanning tree can be more efficiently represented in SAT by treating it as a rooted tree. In the constructed tree, each vertex denotes a row of *M* (i.e. one isoform) and apparently this tree should have *m* + *k* vertices and at most *m* + *k* levels. Otherwise, the problem is infeasible. The high-level idea of the SAT formulation is trying to put each vertex, including both given and missing ones, to a certain level of the tree and construct their parent-child relationship. It is worth noting that such a parent-child relationship is solely for the convenience of construction, it does not indicate the direction of evolution – recall that our model is an undirected graph that primarily concerns about the presence/absence of isoforms, no effort has been made to infer the actual evolution trajectory.

We now provide the implementation details for the above idea. Recall that an SAT formulation consists of a set of boolean/binary variables and a conjunction of clauses where each clause is a disjunction of literals (boolean variables or their negations). We first introduce boolean variable *D*_*i,j*_ to denote whether an edge exists between vertex *i* and vertex *j, i /*= *j*. So *D*_*i,j*_ is True if and only if the *i*-th isoform is derivable from the *j*-th isoform via exactly one simple AS event. Let a helper binary variable *d*_*i,j,k*_ denote the number of (extra) event to convert *M*_*i,k*_ from *M*_*j,k*_ (*i ≠ j*), i.e. flipping the bit of the *k*-th region. Since we only permit one AS event between direct parent-child isoforms, *D*_*i,j*_ is set to True if and only if exactly one variable in *{d*_*i,j,k*_ | 1 *≤ k ≤ n}* is True. Enforcing the condition “exactly one variable in a set must be True” can be implemented as SAT clauses detailed in Suppl. Note S2.

Consider a simplified case when all exons are represented by exactly one region, *i*.*e*. no partial-exons. Then *d*_*i,j,k*_ is set to True if and only if *M*_*i,k*_ ≠ *M*_*j,k*_. However, when partial-exon exists, skipping multiple consecutive partial-exons is also regarded as one event because it takes the same number of splicing to skip one exon or multiple consecutive partial-exons (Fig 1E). Thus, we set *d*_*i,j,k*_ to True, if and only if both conditions are true: (1) *M*_*i,k*_ *M*_*j,k*_; and (2) *M*_*i,k*_ ≠ *M*_*i,k−*1_ or *M*_*j,k*_ ≠ *M*_*j,k−*1_, 2 *≤ k ≤ n*; (second condition not required when *k* = 1). Otherwise the difference between *M*_*i,k*_ and *M*_*j,k*_ has been compensated at or before position *k −* 1, so the penalty should not be double-counted. Those two conditions can be modeled by clauses in Suppl. Note S3.

After properly representing edges with *D*_*i,j*_, we can fit vertices into a tree. Let boolean variable *L*_*i,j*_ denote whether vertex *i* is on level *j* of this tree. *L*_*i,j*_ should satisfy the following constraints: First, a vertex appears exactly once in the tree, which means for the *i*-th isoform, exactly one of the variables in *{L*_*i,j*_ | *∀j}* is set to True. Second, exactly one vertex, *i*.*e*. the root, is on level 1, namely, exactly one variable in *{L*_*i*,1_ | *∀i}* is True. Both require the constraint “exactly one variable in a set is True”, which again can be modeled with the approach in Suppl. Note S2.

Next, we add constraints governing the spanning tree edges. The idea is that if a vertex *i* is present at level *g* (*g ≥* 2), then there must exist a node *j* on level *g −* 1 that has an edge connecting to vertex *i*, namely, *D*_*i,j*_ is True. Let binary variable *C*_*i,j,g*_ denote if vertex *i* is on level *g* of the tree and is preceded by vertex *j* on level *g −* 1 through one simple AS event of edge *D*_*i,j*_. Therefore, *C*_*i,j,g*_ can only be set to True if all three variables *D*_*i,j*_, *L*_*i,g*_ and *L*_*j,g−*1_ are True. However, the reverse direction does not always hold because *D*_*i,j*_ may be true for different pairs of vertices *i* and *j*. Intuitively, a vertex *i* can have multiple potential parents in the graph, but we only choose one in the constructed spanning tree. These constraints can be modeled by 3 SAT clauses:

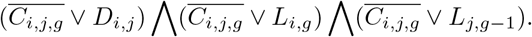

Last, every vertex *i* must be either the root vertex in the spanning tree or located on level *≥* 2. So we have the following constraints for each *i*: exactly one variable from the set *{L*_*i*,1_*}* ∪ *{C*_*i,j,g*_ | *g ≥* 2, *j /*= *i}* is True. Again, Suppl. Note S2 models these constraints.

TENNIS implements the SAT formulation via the pySAT interface [8] and solves the problems using the Glucose SAT solver [4]. We configure it to time-out after 15 minutes to balance computational efficiency and accuracy.

## 3 Results

### 3.1 Most transcript groups satisfy the AS evolution model

In a well-annotated transcriptome, we expect most transcript groups to satisfy our evolutionary model. To verify, we analyzed 7 transcriptome annotations from 6 model species: human (GRCh38 RefSeq and GENCODE), mouse (GRCm39), drosophila (dm6), zebrafish (GRCz11), maize (Zm-B73-REFERENCE-NAM-5.0) and Arabidopsis (TAIR10). For each transcriptome, we first partition all multi-exon transcripts into transcript groups, see Section 2.1. TENNIS is applied to all transcript groups, and according to the outcomes, they are partitioned into 7 categories: 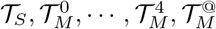.

The statistics are reported in Table 1. It is found that 70%-87% of the transcript groups have just one (multi-exon) transcript. Since different transcript groups have distinct TSS or TTS, this observation aligns well with previous studies that TSS and TTS are the major source of transcriptome diversity [2,24]. Interestingly, human RefSeq has the lowest single-transcript group rate (70%) while human GENCODE has the highest single-transcript group rate (87%). We note that GENCODE has many more transcript groups than Refseq (145202 vs. 90429) but fewer of them are multi-transcript groups (26884 vs. 19425). This indicates GENCODE annotated more genes and alternative TSS/TTS isoforms but fewer AS isoforms per gene.

Among the transcript groups with multiple transcripts (i.e., *𝒯*_*M*_), majority of them (78%–88%, except human/GENCODE) satisfy our model (i.e., in 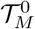), proving the rationality of this model. Human GENCODE is an outlier, with 62% transcript groups ending up in 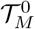. This might be due to a combination that GENCODE over-annotated some transcripts with alternative TSS/TTS isoforms and that some transcript groups are incomplete. Among transcript groups that do not satisfy our model, the majority of them are in 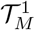, i.e., for most groups, only one isoform is required so as to make it complete. Lastly, approximately only 1% transcript groups requires more than 4 transcripts to meet our model or timed-out in 15 minutes for 6 out of the 7 annotations (it is 3% groups for human GENCODE). This suggests that the parameters of the 15-minute time-out threshold and the maximum number of 4 missing isoforms serves as a sufficient balance between completeness and efficiency for the great majority of transcript groups.

### 3.2 TENNIS-predicted isoforms are validated by long-reads RNA-seq data

It is of great interest to testify whether TENNIS is able to predict correct novel isoforms. Since TENNIS predicts novel/missing isoforms from the reference transcriptome annotations without additional input, if those isoforms can be cross-validated by other data sources such as RNA-seq or external databases, then they are likely to be true positives. In this way, we demonstrate the accuracy and applicability of TENNIS.

We choose the drosophila transcriptome as an example, which is relatively small and well-studied. We retrieved an assembly of high-depth long-read RNA-seq data from a previously published dataset (ref [2]). We used GffCompare [21] to compare the predicted isoforms from TENNIS against this assembly. GffCompare considers two multi-exon isoforms as the same if they have the same intron-chain, which is a widely accepted practice. A TENNIS-predicted novel isoform is considered “supported” if it shares the same intron chain as a transcript from a different source. Otherwise, the prediction is considered “unsupported”. In this experiment, we consider “supported” as true-positive and “unsupported” as false-positive, hence, the count of “supported” predictions as being proportional to the actual recall, and the frequency of “supported” predictions as proportional to the true precision. We also set-up a baseline comparison through randomized approaches. Specifically, recombinations of alternative exons were randomly chosen and coupled with constitutive exons for each group in 𝒯_*M*_. We developed two random baselines. In the first one, referred to as “Rand1”, 1 isoform per transcript group in 𝒯_*M*_ is randomly generated; in the second one, termed “Rand*X*”, *k* novel isoform per group in 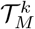 were produced, where the the value of *k* is obtained by TENNIS.

Here, we present a precision-recall plot of TENNIS, Rand1, and Rand*X* (Fig. 2A). Both randomized baselines have lower precisions and lower recalls than TENNIS. The Rand1 and Rand*X* baseline predictions have only 149 and 171 supported isoforms and 23.2% and 18.3% support rates, while TENNIS has 649/691 supported isoforms at the same precision level of Rand1/Rand*X* and support rates of 39.84%/39.86% at the same recall level of Rand1/Rand*X*. Recall that the PctIn (Percentage In) level of a predicted isoform is defined as the number of SAT solutions containing this isoform divided by the total number of solutions for that transcript group. A higher PctIn level indicates a higher confidence that the predicted isoform indeed is missing from the transcript group. At the PctIn level of 0.5 and 0.33, TENNIS reported respectively 41.4%/203 and 30.8%/447 support rate/number (circled points in Fig. 2). Additionally, at the two extremes, TENNIS reported a support rate of 50% for the intersection of all potential solutions and has a support number of 693 for the union of all potential solutions. These observations demonstrated that novel transcripts that have a higher chance of being from the evolution trajectory are more likely to be true positives, which consolidates the evolutionary model of TENNIS.

**Fig. 2:**
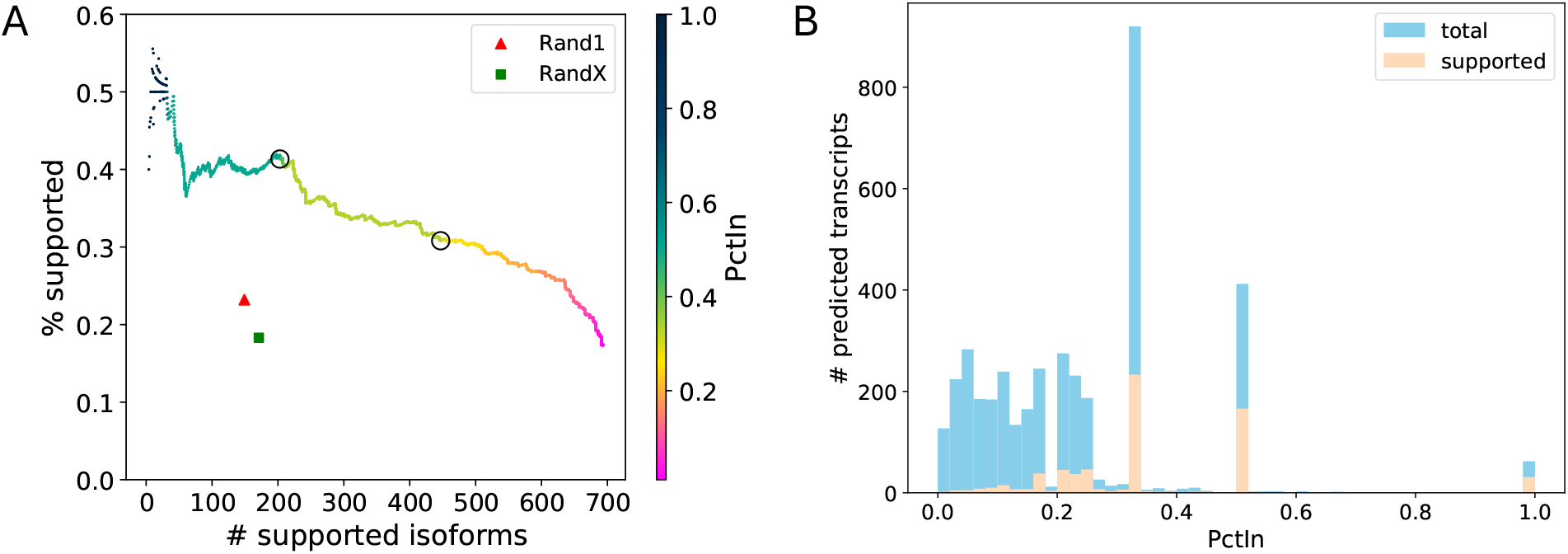
TENNIS augmentations on the Drosophila transcriptome dm6. (A) Precision-recall for transcript predictions sorted by descending PctIn order. The color gradient indicates PctIn values, with circles highlighting critical thresholds (PctIn = 0.5, 0.333). (B) Histogram of PctIn values for predicted transcripts. Three local peaks were observed at 0.333, 0.5 and 1.0 PctIn.

Although the PctIn values indeed range from 0 to 1, their distribution displays significant skewness with discrete peaks occurring at 0.333, 0.50, and a smaller local peak at 1.0 (Fig. 2B). Transcripts with PctIn values of 0.333 (resp. 0.50 or 1.0) are possibly from a transcript group *T* where each of them has three (resp. two or one) optimal solutions from SAT. Note this concept is different from 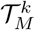 which describes the number of missing isoforms. In other words, a transcript group *T* may need only one isoform to form a connected graph (thus, in 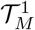), but may have two possible optimal configurations for this isoform by SAT. Correspondingly, transcripts with lower PctIn values are from a transcript group *T* with more optimal SAT solutions. Apparently, the latter group is harder to solve, and predicted isoforms from such groups are less favorable. Transcripts with PctIn values of 0.333 (resp. 0.50 or 1.0) have a precision of 25% (resp. 40% or 51%), much higher than that of transcripts with lower PctIn values (9.6%). Therefore, we show that PctIn values of 0.5 and 0.333 can generally serve as two good thresholds for filtering TENNIS predictions.

It is noteworthy that not all genes or transcripts are expressed. Also, not all that expressed are sequenced. Hence, using assemblies from real RNA-seq as a ground truth tends to underestimate the total number of true-positive genes and/or transcripts. In other words, transcript predictions un-supported by an assembly may be false-positive predictions or due to being unexpressed/unsequenced in the experiments. To estimate the coverage of genes in our “ground-truth” (namely, the assembly from ref [2]), we compared it with dm6 annotations, in addition to TENNIS outputs. GffCompare reported this long-read assembly overlaps with only 54.0% loci (based on exon overlapping [21]) in dm6 annotation and 63.0% loci in TENNIS. Hence, the number of true positives is most likely underestimated for TENNIS to a noticeable level.

### 3.3 TENNIS accurately retrieves isoforms in a removal simulation

To further validate TENNIS’s ability to detect missing transcripts, we conducted a simulation using a removal and retrieval approach. From genes containing three or more annotated isoforms, we randomly removed one isoform. The removed one cannot be the shortest isoform and the removal of it should not reduce the total number of exons (i.e. the exon spliced out in all other isoforms) in the group, so that retrieval of this isoform is not impossible. This experimental design aimed to assess both the precision and recall of TENNIS in identifying missing transcripts. GffCompare was used for evaluation and the removed transcripts are regarded as ground truth.

A total of 796 multi-isoform groups were used for this removal simulation. TENNIS classified the 796 *𝒯*_*M*_ groups to 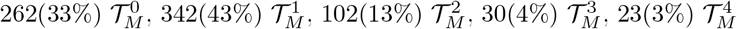 and 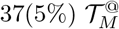 groups. The percentages of all 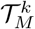 classes increased, compared to Table 1. This is natural since we removed one isoform from each group. The presence of 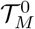 groups indicates that some groups have a more “connected” graph and that not all non-terminal vertices are cut vertex. Besides, those 796 groups do not necessarily satisfy the evolution model prior to the removal. Therefore the missing isoform identification problem is further entangled.

TENNIS achieved high precision and recall, which is considerably better than randomized approaches, in this simulated removal-and-retrieval experiment (Fig. 3A). At PctIn values of 0.5 or 0.333, TENNIS has a precision of 40.6% or 27.9% and a recall of 202 or 329. The precision and recall for Rand1 are 19.5% and 97, while those for Rand*X* are 14.7% and 107. TENNIS has a precision of approximately 45% or a recall of 375 isoforms after controlling recall or precision at a similar level. The above numbers are substantially higher than those for the two randomized baselines.

**Fig. 3:**
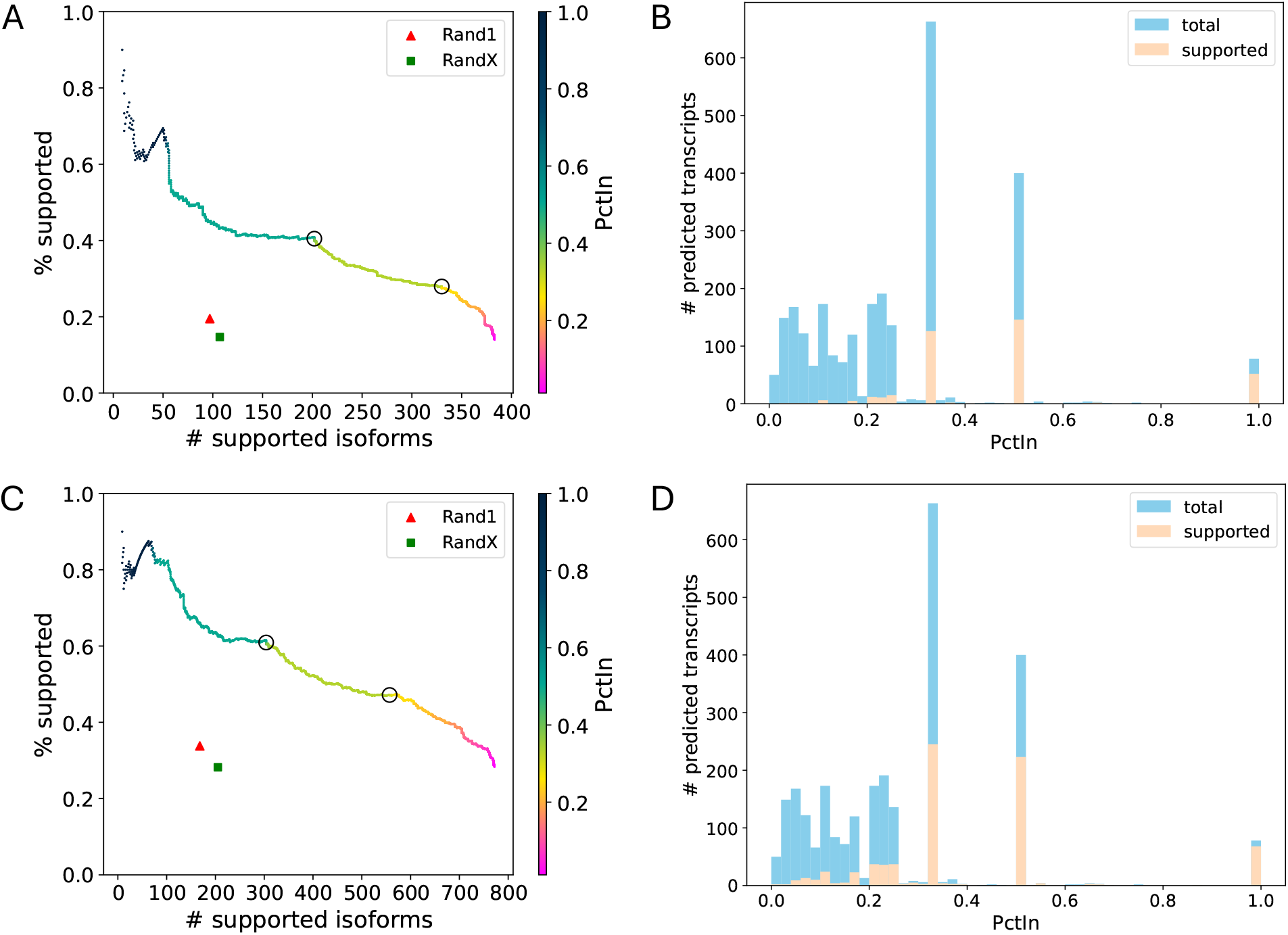
(A) Precision-recall for transcript predictions sorted by descending PctIn order, validated against exactly removed isoforms. The color gradient indicates PctIn values, with circles highlighting critical thresholds (PctIn = 0.5, 0.333). (B) Histogram of PctIn values for predicted transcripts validated against exactly removed isoforms. (C) Precision-recall analysis using the same methodology and color scheme as panel A, validated against combined ground truth (exactly removed isoforms + long-read RNA-seq assembly). (D) Histogram of PctIn values for predicted transcripts validated against combined ground truth.

Considering the presence of multiple solutions and the potential incompleteness of annotations, we also evaluated the predictions using the combined ground truth, i.e. union of removed transcripts and the RNA-seq assembly (Fig. 3C). At a PctIn level of 0.5 (resp. 0.333), TENNIS successfully predicts 304 (resp. 556) supported isoforms with a support rate of 61.0% (resp. 47.1%). The baseline approaches Rand1 and Rand*X* respectively only identified 168 and 205 supported isoforms with support rates of 33.8% and 28.2%. In contrast, TENNIS has more than doubled precisions and approximately 3.5-4.5 times supported isoforms under the same recall or precision of the two baselines, showing remarkable improvements.

The distribution of PctIn values mirrors the pattern observed in Seciton 2, exhibiting local peaks at 0.33, 0.5, and 1.0 (Fig. 3B and D). The precisions of isoforms with those PctIn values are 19%, 37%, 67% if validated by exact removed isoforms, and 37%, 56%, 87% if validated by combined ground truth.

## 4 Conclusion and Discussion

A comprehensive transcriptome annotation is essential for many bioinformatic and biomedical studies. While significant resources have been invested in improving these annotations through the invention of new methods, pipelines, and manual curations, the great majority of them, if not all, are data-driven [14,1,22,17]. Little attention has been paid to modeling the annotated isoforms particularly through an evolution perspective. We fill this critical gap with TENNIS, an evolutionary model for characterizing annotated transcripts, together with an algorithm that infers missing isoforms in an annotation. The model of TENNIS is simple: isoforms in a transcript group are connected in a single component using the four basic AS events, should no isoform missing. When this condition is not satisfied, TENNIS seeks the minimum number of isoforms to make them connected, using a novel SAT formulation that guarantees to find all optimal solutions.

We analyzed seven transcriptome annotations of model organisms using TENNIS. It was shown that the majority of transcript groups are single-isoform, accounting for about 80% of all multi-exon groups. The evolution model is satisfied by 62%-88% of transcript groups in various species’ annotations, consolidating the propositions of our model. We also evaluated the validity of TENNIS isoform predictions by comparing them with a long-read RNA-seq assembly and through a simulation experiment of removal and retrieval. In both settings, we demonstrated that TENNIS dramatically outperformed two randomized baseline methods. After controlling the same level of recall or precision, TENNIS showed approximately a 70%-200% increase of precision and 250%-330% increase of recall over the baselines in the experiments. Furthermore, the analysis revealed that if an isoform appears in multiple optimal solutions with a higher percentage (PctIn), then it is more likely to represent a true isoform. The PctIn metric can thus serve as an effective criterion for filtering predicted isoforms.

The assumption made by TENNIS is minimal, yet demonstrates strong prediction capability in identifying missing isoforms. It therefore holds great potential to model complex evolutionary trajectories. A promising future enhancement for TENNIS is the integration of additional prior knowledge and features related to (AS) events, such as the lengths and sequences of introns/exons and their splicing patterns. Previous studies have established that constitutive exons typically exhibit greater lengths and are flanked by shorter introns, while alternatively spliced exons are more likely to be shorter and accompanied by longer introns [3,13,11]. Also, ref [13] unveiled the positive correlation between expression level and evolutionarily conserved transcripts, which are often ancestral. As a mathematical model to characterize observed isoforms, TENNIS provides valuable insights for various studies including alternative splicing mechanisms, comparative transcriptomics, and phylo-transcriptomics studies.

## 5 Code Availability

TENNIS is freely available at https://github.com/Shao-Group/tennis. Scripts and documentation for reproducing the experiments in this manuscript are also available at https://github.com/Shao-Group/tennis-test.

## 6 Acknowledgments

The authors thank Tasfia Zahin for help in data collection and processing. This work is supported by the US National Science Foundation (2145171 to M.S.) and by the US National Institutes of Health (R01HG011065 to M.S.).

