## supplemental materials for "Augmenting Transcriptome Annotations through the Lens of Splicing Evolution"

### Suppl. Note S1

Let Boolean variable $E$ denote the equality of two Boolean variables $X$ and $Y$. That means $E$ must be set to True if and only if $X = Y$, and $E$ must be set to False if and only if $X \neq Y$. This is constrained via four clauses:

$$X \vee Y \vee E$$
$$\overline{X} \vee \overline{Y} \vee E$$
$$X \vee \overline{Y} \vee \overline{E}$$
$$\overline{X} \vee Y \vee \overline{E}$$

Similarly, inequality can also be expressed in SAT clauses. Let Boolean variable $E'$ denote the inequality of two Boolean variables $X$ and $Y$. That means $E'$ must be set to False if and only if $X = Y$, and $E'$ must be set to True if and only if $X \neq Y$. This is constrained via four clauses:

$$X \vee Y \vee \overline{E'}$$
$$\overline{X} \vee \overline{Y} \vee \overline{E'}$$
$$X \vee \overline{Y} \vee E'$$
$$\overline{X} \vee Y \vee E'$$

.

### Suppl. Note S2

Given a set of Boolean variables $\{v_1, v_2, \cdots, v_n\}$, we want to constrain that exactly one variable is True and all the other variables must be False. One way to formulate this constraint in conjunctive normal form (CNF) is through pairwise cardinality encoding [1].

First, at least one of $\{v_1, v_2, \cdots, v_n\}$ is True:

$$\bigvee_{i=1}^{n} v_i$$

And, second, no two of $v_i$ are both True:

$$\overline{v_i} \vee \overline{v_j}, \forall i \neq j$$

### Suppl. Note S3

Given an $m \times n$ matrix $M$, we want to compute the number of (extra) event to convert $M_{i,k}$ from $M_{j,k}$ at region $k$. Apparently, either 0 or 1 event is needed to convert one exon to intron or the contrary. Let this number be denoted by the helper binary variable $d_{i,j,k}$.

Then, $d_{i,j,k}$ is set to True, if and only if both conditions are true: (1) $M_{i,k} \neq M_{j,k}$; and (2) $M_{i,k} \neq M_{i,k-1}$ or $M_{j,k} \neq M_{j,k-1}$ $(2 \leq k \leq n)$. The second condition is not required when $k = 1$. To simplify the representation and reduce the number of clauses, let binary helper variable $Eq_{i,k}$ denote whether $M_{i,k} = M_{i,k-1}$ $(k \geq 2)$ using clauses in Suppl. Note S1.

Afterward, we need all of the following clauses to be satisfied.

First, $d_{i,j,k}$ is set to False, if $M_{i,k} = M_{j,k}$:

$$M_{i,k} \vee M_{j,k} \vee \overline{d_{i,j,k}}$$

$$\overline{M_{i,k}} \vee \overline{M_{j,k}} \vee \overline{d_{i,j,k}}$$

Second, $d_{i,j,k}$ is set to False, if $Eq_{i,k}$ and $Eq_{j,k}$ are both True:

$$\overline{Eq_{i,k}} \vee \overline{Eq_{j,k}} \vee \overline{d_{i,j,k}}$$

Third, $d_{i,j,k}$ is set to True, if (1) $M_{i,k} \neq M_{j,k}$ and (2) $Eq_{i,k}$ and $Eq_{j,k}$ are not both True:

$$M_{i,k} \vee \overline{M_{j,k}} \vee Eq_{i,k} \vee d_{i,j,k}$$

$$M_{i,k} \vee \overline{M_{j,k}} \vee Eq_{j,k} \vee d_{i,j,k}$$

$$\overline{M_{i,k}} \vee M_{j,k} \vee Eq_{i,k} \vee d_{i,j,k}$$

$$\overline{M_{i,k}} \vee M_{j,k} \vee Eq_{j,k} \vee d_{i,j,k}$$
